# SOAR elucidates disease mechanisms and empowers drug discovery through spatial transcriptomics

**DOI:** 10.1101/2022.04.17.488596

**Authors:** Yiming Li, Saya Dennis, Meghan R. Hutch, Yanyi Ding, Yadi Zhou, Yawei Li, Maalavika Pillai, Sanaz Ghotbaldini, Mario Alberto Garcia, Mia S. Broad, Chengsheng Mao, Feixiong Cheng, Zexian Zeng, Yuan Luo

**Affiliations:** Department of Preventive Medicine, Northwestern University, USA; Genomic Medicine Institute, Lerner Research Institute, Cleveland Clinic, USA; Department of Molecular Medicine, Cleveland Clinic Lerner College of Medicine, Case Western Reserve University, Cleveland, OH, USA; Case Comprehensive Cancer Center, Case Western Reserve University, Cleveland, OH, USA; Center for Quantitative Biology, Academy for Advanced Interdisciplinary Studies, Peking University, Beijing, China; Peking-Tsinghua Center for Life Sciences, Academy for Advanced Interdisciplinary Studies, Peking University, Beijing, China

## Abstract

Spatial transcriptomics provides researchers with a better understanding of gene expression within the tissue context. Although large volumes of spatial transcriptomics data have been generated, the lack of systematic curation and analysis makes data reuse challenging. Herein, we present Spatial transcriptOmics Analysis Resource (SOAR), a resource with an extensive, systematically compiled collection of spatial transcriptomics data across tissues, organs, and pathological conditions. SOAR is a comprehensive database with uniformly processed and annotated samples, facilitating future benchmark studies and method development. SOAR also offers multi-pronged analysis capability, including an integrative approach toward drug discovery that allows for efficient exploration of novel and targeted therapeutic uses for existing compounds.

## Main

Spatially resolved transcriptomics preserves the spatial organization of capture locations, which allows researchers to study tissue functions and disease pathology in the morphological context^1^. In recent years, different spatial transcriptomics technologies have been widely applied, leading to discoveries in cancer research, neuroscience, and developmental biology. To facilitate data exploration and meta-analysis in spatial transcriptomics research, we present Spatial transcriptOmics Analysis Resource (SOAR; https://soar.fsm.northwestern.edu/), a public database with an extensive collection of spatial transcriptomics data, analysis functions, and interactive visualization features. SOAR hosts an interactive web interface for users to analyze and visualize spatial gene expressions in different tissue samples. SOAR also offers functions for users to analyze spatial variability and cell-cell interactions, as well as perform drug discovery/repurposing across datasets, allowing systematic comparisons across samples, cell types, and disease types. Various data resources for spatial transcriptomics currently exists, such as SpatialDB^2^, STOmicsDB^3^, SPASCER^4^, SODB^5^, Aquila^6^, Museum of Spatial Transcriptomics^7^, spatialLIBD^8^, STAR-FINDer^9^, Amyotrophic Lateral Sclerosis Spinal Cord Atlas^10^, and SORC^11^.However, none of these resources possesses a comparable size and range of functionalities when compared to SOAR. SpatialDB^2^ is a manually curated spatial transcriptomics database with 305 samples. It provides spatial expression visualization and pre-computed results for sample-wise spatially variable gene identification. STOmicsDB^3^ hosts 1,033 spatial transcriptomics samples, of which 463 samples are processed and have spatial coordinates. Users can perform sample-wise clustering and spatial variability analysis on the processed datasets using STOmicsDB. SPASCER^4^ is a spatial transcriptomics database providing clustering, spatial variability, pathway, gene regulatory network, and neighborhood-based cell-cell interaction analyses. Out of all the 1,082 samples on SPASCER, 1,030 samples have spatial transcriptomics data analysis results. SODB^5^ is a spatial transcriptomics database with 1,701 samples, and users may use SODB to perform spatial expression visualization, clustering, and spatial variability analysis. Aquila^6^ is a platform allowing users to explore spatial transcriptomics data through spatial expression visualization, clustering, spatial variability, marker co-localization, and neighborhood-based cell-cell interaction analysis. Aquila^6^ currently hosts 1,397 samples with data analysis results. Museum of Spatial Transcriptomics^7^ is an annotated list of spatial transcriptomics literature with links to publications and their datasets. Multiple data atlases also exist, including spatialLIBD^8^, STAR-FINDer^9^, Amyotrophic Lateral Sclerosis Spinal Cord Atlas^10^, and SORC^11^. These atlases provide interfaces for visualizing spatial expressions and data analysis, but their scopes are limited to a particular tissue type or disease category (brain, intestine, spinal cord, and cancer, respectively). To our best knowledge, SOAR is the largest spatial transcriptomics database that offers unified data access, exploration, and analysis functions across multiple tissue types, and the first spatial transcriptomics database providing drug discovery results for different disorders. SOAR hosts a substantial collection of systematically curated and processed datasets (2,785 samples), which can facilitate future benchmark studies and method development. SOAR also provides user-friendly functions including interactive visualization of spatial gene expression and clustering results, exploration of spatial variability, analysis of neighborhood-based and distance-based cell-cell interactions, screening of potential drugs, and meta-analyses across datasets (**Fig. 1a**, Supplementary Table 1). In summary, SOAR is a comprehensive resource that can improve the understanding of gene functions and disease pathology in the tissue context as well as facilitate therapeutics discovery.

**Fig. 1:**
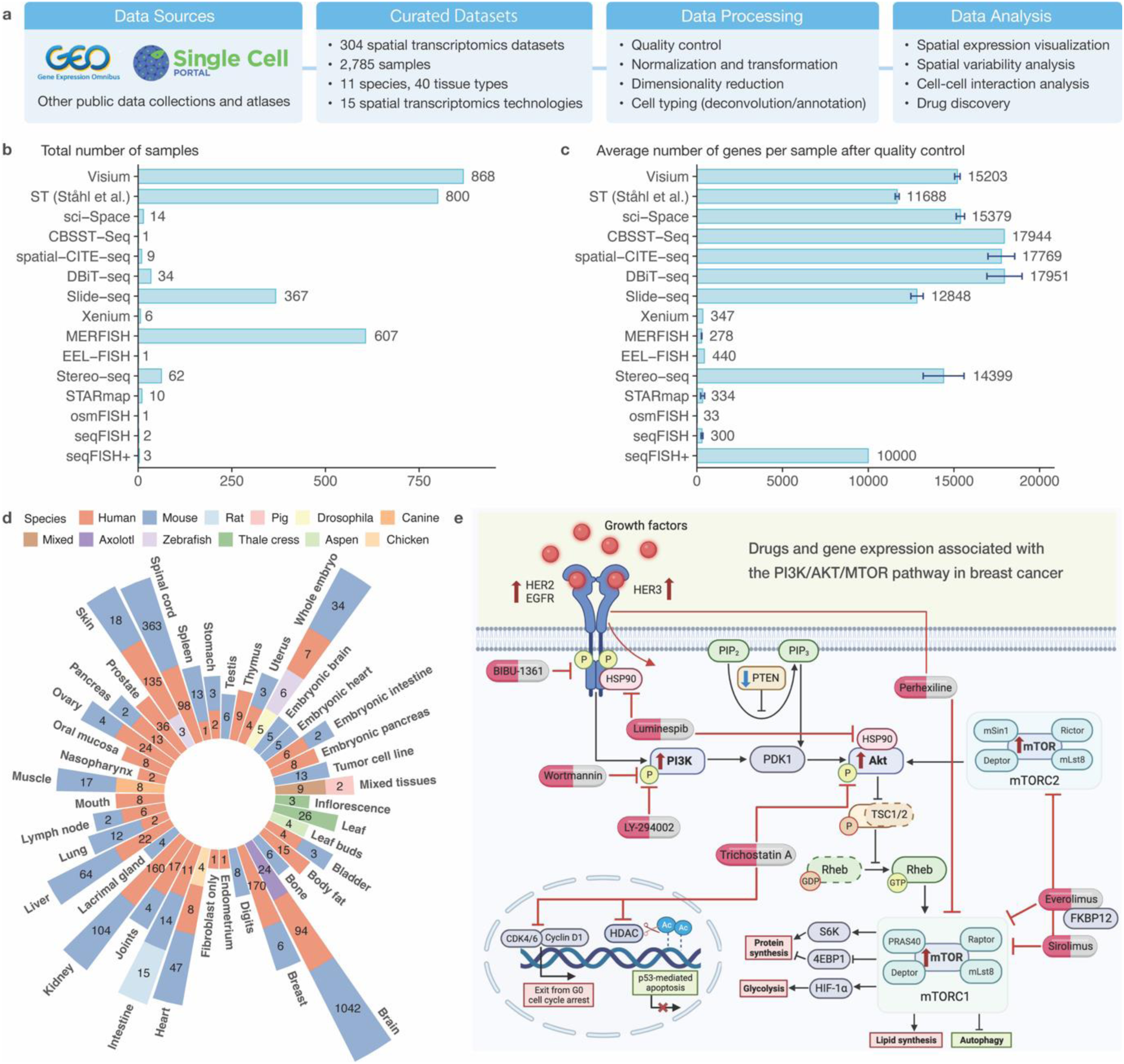
Overview of SOAR and drugs targeting genes in the PI3K/Akt/mTOR pathway in breast cancer. Datasets hosted on SOAR were curated from public domains and processed using a standardized workflow. SOAR also provides interactive interfaces for users to visualize spatial gene expressions, evaluate the spatial variability of genes, assess cell-cell interactions, and conduct drug discovery. **b-c**, Statistics of data from different spatial transcriptomics technologies in SOAR. The 95% confidence intervals for the means are plotted as error bars. **d**, Number of samples of each species and tissue type combination. SOAR contains 2,785 spatial transcriptomics samples from 11 different species across 40 tissue types. **e**, The drug discovery function of SOAR identifies various compounds that target the PI3K/Akt/mTOR growth and proliferation pathway and show strong suppression effects toward differentially expressed genes in malignant cells.

We systematically curated 2,785 samples across 11 species and 40 tissue types from 304 datasets (**Fig. 1d**, Supplementary Table 2). The curated datasets were generated by 15 different spatial transcriptomics technologies (**Fig. 1b-c**). The datasets were reviewed, annotated, and pre-processed by a standardized workflow (Methods). In SOAR’s data browser, users could browse the curated datasets to pinpoint samples of interest using metadata-based filters and visualize spatial expressions interactively (**Fig. 2a**). More importantly, the processed data, including normalized expressions, images, coordinates, and phenotypic data, are available to download for all samples. In addition to providing data access and visualization, SOAR could help users evaluate gene spatial variability, investigate cell-cell interactions, and discover repurposable drugs across various datasets.

**Fig. 2:**
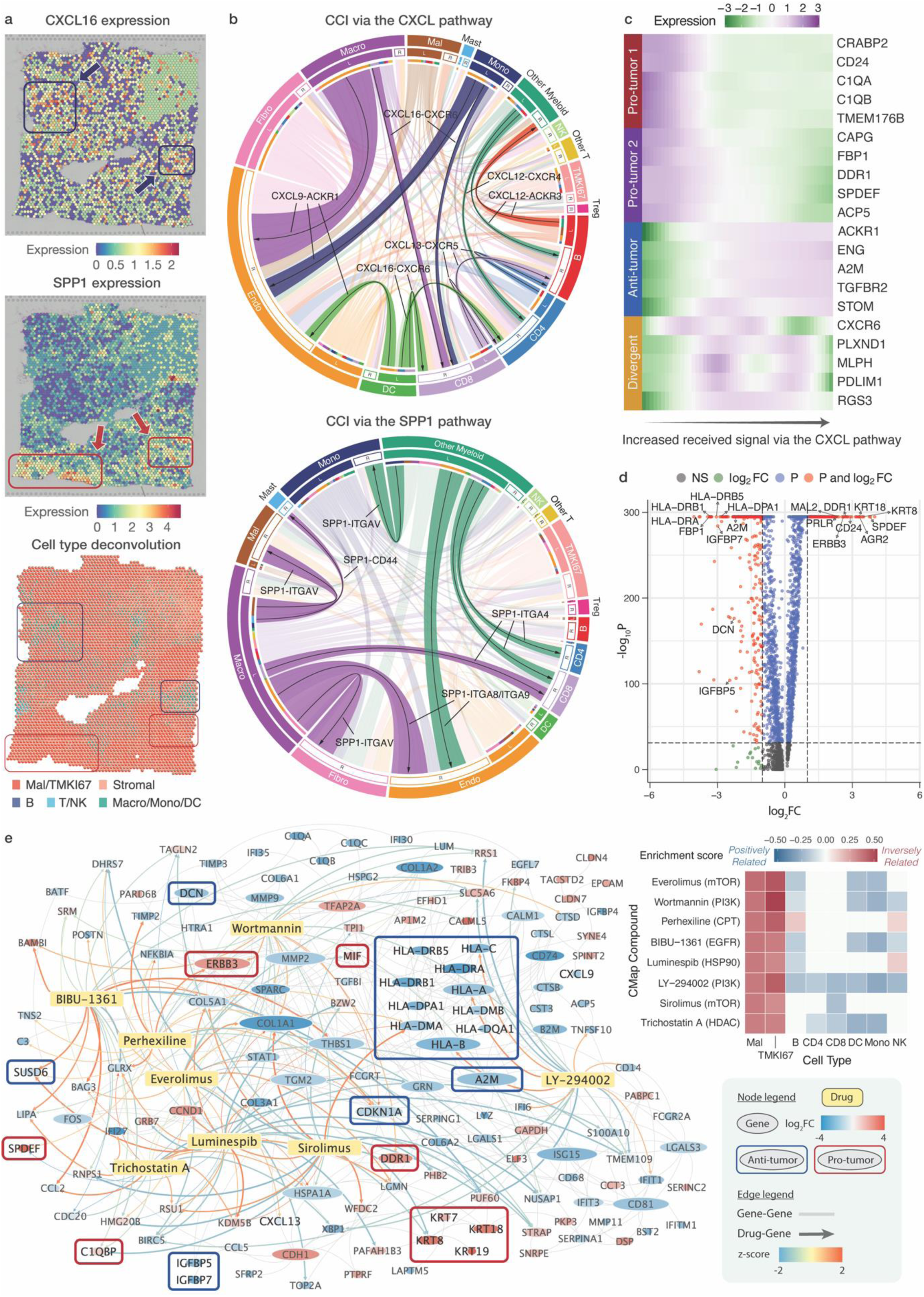
Explore cell-cell interactions and perform drug discovery in cancer spatial transcriptomics data in SOAR. Users can use SOAR to visualize (**a**) spatial gene expressions and cell type deconvolution results in a sample of interest. To study cell-cell interactions, users can explore (**b**) distance-based cell-cell interactions and (**c**) signaling differential genes. To conduct drug discovery, users can visualize (**d**) differentially expressed genes in a studied cell type and (**e**) protein-protein interactions and drug perturbations. **a**, Spatial gene expressions of CXCL16 and SPP1 in a breast cancer sample. The arrows indicate distinct regions of elevated expression of the genes. **b**, Pan-cancer short-distance (500 μm) cell-cell interactions among different cell types through the CXCL and SPP1 pathways. The top ligand-receptor pairs are highlighted. **c**, Heatmap of selected genes differentially expressed with respect to the amount of received signal through the CXCL pathway in a breast cancer sample. **d**, Volcano plot of the genes differentially expressed in malignant cells in a breast cancer sample. Known tumor-promoting and tumor-suppressing genes are highlighted. The absolute log fold change threshold is set at 1, and the adjusted p-value threshold is set at 10^-32^. **e**, Protein-protein interaction and drug perturbation network in a breast cancer sample. The gene nodes are colored by their log fold change in malignant cells compared with other cell types and sized by the total degree. Known anti-tumor and pro-tumor genes are boxed. The edges between drugs and genes are colored and weighted by the expression z-score from Connectivity Map. The enrichment scores of the selected drugs are visualized in a heatmap colored by their enrichment scores with respect to different cell types. CMap, Connectivity Map; DC, dendritic cells; Endo, endothelial cells; Fibro, fibroblasts; log2FC, log-fold change; Macro, macrophages; Mal, malignant cells; Mono, monocytes; NK, natural killer cells; NS, non-significant; Treg, regulatory T cells; P, false-discovery-rate-adjusted p-value; z-score, expression z-score from Connectivity Map.

The spatial patterns of gene expression inform us about the migration of cells, their responses to different tissue environments, and cell-cell interactions^1, 12^. SOAR allows users to evaluate the statistical significance of a gene’s spatial variability in different tissue samples and cell types. Chemokine genes like CXCL16 are known to be associated with immune activation and the prognosis of cancer^13, 14^, and their expression levels were found to have significant spatial pattern variation in cancer samples^12, 15^. SPP1 (secreted phosphoprotein 1) has been shown to be overexpressed in different types of tumors^16, 17^ and promote tumor progression and metastasis^18, 19^. Therefore, as a case study, we visualized the spatial variability of CXCL16 and SPP1 in our curated breast cancer samples (Supplementary Fig. 1). Our results suggested significant spatial expression variation of both genes for most breast cancer samples (Supplementary Fig. 1), indicating that the genes have high activity in certain microenvironments only, as opposed to house-keeping genes which tend to be highly expressed throughout the tissue without any spatial pattern. More specifically, in different samples, the two genes have significant (adjusted p-value < 0.05) spatial variability more frequently in macrophages compared with other cell types, indicating cell type specificity in their roles in the tumor immune microenvironment (Supplementary Fig. 1). In addition, our spatial expression visualization revealed that CXCL16 and SPP1 colocalize with different cell types (**Fig. 2a**). CXCL16 was highly expressed in areas enriched with T and natural killer (NK) cells (**Fig. 2a**). This can be attributed to the fact that CXCL16 expression could be induced by interferon-γ that was primarily secreted by T and NK cells^20, 21^. In addition, areas with low CXCL16 expression and high malignant cell proportion exhibited elevated levels of SPP1 (**Fig. 2a**). This spatial pattern aligns with previous findings that tumor progression can induce SPP1 expression in macrophages^22^. Our results indicate that CXCL16 and SPP1 are likely associated with pro-inflammatory and pro-tumor responses, respectively, which aligns with previous findings^12, 13, 16, 23^. Altogether, these results demonstrate that even without histological annotation, SOAR could identify biologically relevant genes with localized expression patterns.

Cell-cell interactions (CCIs) may occur over a long distance through the secretion of chemokines^24^. With SOAR, users could explore if the expression of a specific gene correlates with the presence of nearby cell types (Supplementary Fig. 3). In addition, SOAR also enables users to investigate far-reaching but biologically meaningful CCI signals (**Fig. 2b-c**, Supplementary Fig. 2). Our spatial variability analysis shed light on the divergent roles of CXCL16 and SPP1 in the tumor immune microenvironment (TME). To delve deeper, we evaluated cancer sample CCIs related to chemokine signaling (mainly the CXCL16-CXCR6 ligand-receptor pair) and the SPP1 gene (mainly the SPP1-ITGAV interaction). We found that while CXCL16-CXCR6 activated immune responses, SPP1-ITGAV may lead to tumor growth (**Fig. 2b**). In previous studies, CXCR6+ CD8 T cells were found to possess stronger anti-tumor ability and showed an enhanced response to PD-1 inhibitors^25^. This finding was confirmed in spatial transcriptomics data, as our analysis showed that CXCL16+ dendritic cells (DCs) and myeloid cells interacted with CXCR6+ CD8 T cells, signifying a potential synergistic immune response against tumor (**Fig. 2b**). Other chemokine interactions that also helped recruit immune cells^26^ and increase lymphocyte infiltration^27^ included CXCL13+ T cells and CXCR5+ B/T cells (**Fig. 2b**). Furthermore, our differential expression analysis showed that more signal from CXCL16-CXCR6 and other chemokine-related CCIs increased the expressions of tumor-suppressing genes like ACKR1^28, 29^, A2M^30^, and TGFBR2^31, 32^, while decreased the expressions of tumor-promoting genes like CD24^33^, C1QA/C1QB^34, 35^, and DDR1^36^ (**Fig. 2c**). Our neighborhood-based CCI analysis also demonstrated the crosstalk between tumor and immune cells in mediating immune escape. For example, tumor cells are more likely to express immunosuppressive complement genes C1QA/C1QB when located near macrophages (Supplementary Fig. 3). This result suggested positive interactions between malignant cells and tumor-associated macrophages in evading immune surveillance^35^. On the other hand, the CCIs involving SPP1 occurred more frequently among macrophages and stromal cells rather than macrophages and other immune cells (**Fig. 2b**). For example, SPP1+ macrophages strongly interacted with ITGAV+ endothelial cells, fibroblasts, and malignant cells (**Fig. 2b**). SPP1’s involvement in the integrin pathway has been shown to lead to metastatic seeding^37^ and activation of the Akt pathway^17^. The silencing of ITGAV can in turn inhibit tumor growth^37^. In addition, SPP1+ macrophages also interacted with CD44+ mast cells (**Fig. 2b**). Previous studies showed the SPP1-CD44 interaction could hinder mast cells’ ability to activate T cells in the TME^38, 39^. These results suggest that CCIs involving CXCL16-CXCR6 may lead to immunoactivation whereas those related to SPP1-ITGAV may result in immunosuppression.

To further investigate whether CXCL16/SPP1 macrophage polarity is associated with tumor control and progression mechanisms, we analyzed the correlations between the macrophage CXCL16/SPP1 ratio and the expression of cytotoxicity markers and tumor progression markers. Our results showed that the CXCL16/SPP1 ratio was positively correlated with the infiltration of cytotoxic CD8 T cells, which were annotated by the markers GZMA (granzyme A), NKG7 (natural killer granule protein 7), and CCL5 (chemokine C-C motif ligand 5). In addition, the macrophage CXCL16/SPP1 ratio is negatively correlated with tumor progression markers of CDKN2A (cyclin-dependent kinase inhibitor 2A), LDHA (lactate dehydrogenase A), and HIF1A (hypoxia-inducible factor 1 subunit alpha) (Supplementary Fig. 2b-c). Together, these results suggest that macrophage polarity on the CXCL16/SPP1 axis could characterize the coordination of anti-tumor and pro-tumor pathways in the TME. This aligns with recent findings that a higher CXCL9/SPP1 ratio in macrophages is associated with increased tumor infiltration activities and enhanced cancer patient survival^23^.

Joint analysis of transcriptomics data from SOAR and drug perturbation data can aid efficient drug discovery (Methods). Many of the compounds identified from our drug screen that show strong suppression toward malignant cells target the PI3K/Akt/mTOR pathway (**Fig. 1e**). The PI3K/Akt/mTOR pathway promotes tumor growth, invasion, and endocrine resistance^40–44^. Therefore, much effort has been dedicated to developing drugs that inhibit this pathway in recent years^40–44^. We observed higher expression of PI3K, Akt, and mTOR as well as upstream activators HER2 and HER3 but not the tumor suppressor PTEN in our case study sample. The PI3K/Akt/mTOR pathway is activated by the heterodimerization of HER3 with other receptors following the binding of growth factors, which are overexpressed in the TME. A cascade of phosphorylation activity ultimately activates mTORC1, which upregulates biosynthesis and metabolism and represses autophagy in tumor cells^45–48^, as reflected from the differentially expressed genes (DEGs) of the case study sample.

The eight compounds identified from our drug screen that target the PI3K/Akt/mTOR pathway include sirolimus, trichostatin A, everolimus, LY-294002, wortmannin, BIBU-1361, perhexiline and luminespib. These compounds are highly selective toward cancer cells as they can suppress the expression of DEGs in malignant cells without affecting DEGs in immune cells (**Fig. 2e**). Sirolimus is one such compound that can inhibit mTOR via a three-way interaction with FKBP12^48, 49^. As shown from perturbation results from the Connectivity Map (CMap), sirolimus decreases the expression of glycolysis and pentose phosphate pathway genes that were upregulated by mTORC1 in response to hypoxia from excessive growth^47, 50–52^ (Supplementary Fig. 4e). Likewise, sirolimus also reverses the heightened expression of lipid and sterol synthesis genes that are downstream of mTORC1^46, 50^ (Supplementary Fig. 4e). In addition, sirolimus can also suppress mRNA translation, promote autophagy, and reduce mTORC2 to below the level needed to maintain Akt signaling^46–48, 50, 53, 54^. Sirolimus and its analog everolimus have also shown clinical efficacy in treating various solid tumors^55, 56^. The high ranking of sirolimus and everolimus among CMap compounds in the ability to suppress malignant cells’ DEGs confirms the effectiveness of SOAR’s drug screen function in identifying potent antineoplastic drugs via a data-driven approach. Trichostatin A (TSA) is another drug identified from SOAR’s drug screen that strongly suppresses DEGs of malignant cells. While approved as an antifungal, TSA exhibits potential for re-purposing as a cancer drug due to its roles as a class I histone deacetylase (HDAC) inhibitor and Akt inhibitor. TSA acetylates the tumor suppressor p-53, which activates the transcription of downstream apoptosis genes^57, 58^. It can also induce cell cycle arrest by downregulating cyclin D1 and CDK4 while upregulating their inhibitor p21^59, 60^. Both of these aforementioned perturbations are supported by the observed gene expression changes from CMap following TSA treatment (Supplementary Fig. 4e). In addition, TSA can also inhibit phosphorylation of Akt, restore the role of TSC1/2 as GTPase-activating proteins, and prevent Rheb from activating mTORC1^59, 60^. Moreover, HDAC inhibitors could help overcome PI3K/Akt inhibitor resistance through epigenetic modifications^61^. Like everolimus, HDAC inhibitors like TSA demonstrated high selectivity for cancer cells *in vivo* and have undergone evaluation in multiple clinical trials targeting solid tumors^62–65^. The identification of sirolimus and TSA from SOAR’s drug screen underscores our platform’s ability to identify both established and repurposable drugs at a personalized therapy level. More importantly, SOAR also allows users to explore the mechanisms of action of identified compounds by showing gene perturbation results of these compounds from experiments conducted by CMap.

The drug perturbation network of the compounds that disrupt the PI3K/Akt/mTOR pathway is shown along with the protein-protein interaction (PPI) network of malignant cells from the case study (**Fig. 2e**). Most genes perturbed by the drugs are either directly DEGs of malignant cells or interacting with them, suggesting the ability of these compounds to disrupt dysregulated tumor processes. In addition to the pro-tumor and anti-tumor genes identified from distance-based cell-cell interaction analysis, several other tumorigenesis genes were identified. ERBB3 stands out as one of the most highly upregulated genes that also have many PPIs, which explains why many of the top-ranked compounds from drug screening target the downstream PI3K/Akt pathway. In addition, Class II major histocompatibility complex (MHC) genes are downregulated, which aids immune escape in tumor cells^66, 67^.

Interestingly, the PI3K inhibitor wortmannin and CPT inhibitor perhexiline did not suppress ERBB3 expression. However, this observation is not unexpected as there is a negative feedback loop with PI3K inhibition that upregulates ERBB3 mRNA and promotes the receptor’s heterodimerization^68, 69^. Therefore, combination therapy of PI3K/Akt/mTOR inhibitor and HER2 antibody has shown greater efficacy against tumor cells than PI3K inhibitor alone^70^. LY-294002, sirolimus, and BIBU-1361 upregulate MHC genes HLA-A and HLA-DMA while downregulating MHC inhibitor SUSD6, which helps to improve tumor infiltration by lymphocytes. In addition, both BIBU-1361 and wortmannin show strong upregulation of CDKN1A (p21), which inhibits cyclin*-*CDK-mediated cell cycle progression. Lastly, everolimus downregulates MIF, which promotes angiogenesis in cancer cells and whose inhibition showed growth suppression in various in-vivo models^71, 72^. Analyzing the PPI and gene targets of highly ranked compounds from drug screen can further help users to pinpoint the specific pathways that are aberrant in pathological samples and being reversed by the compounds.

In summary, SOAR hosts a large number of spatial transcriptomics datasets that were systematically processed and annotated. Besides data access and download, SOAR provides interactive analysis functions for visualizing spatial gene expression, evaluating gene spatial variability, studying neighborhood-based and distance-based cell-cell interactions, and identifying potential drugs for treating pathological samples. SOAR will be continuously maintained to function as an open-access platform that offers scalable utility to the biomedical and clinical research communities, enabling researchers to fully leverage the potential of spatial transcriptomics.

## Methods

### Data Collection

We queried the Gene Expression Omnibus (GEO, http://www.ncbi.nlm.nih.gov/geo/) for human and mouse spatial transcriptomics datasets using the keywords “spatial+transcriptomics”, “spatial+transcriptome”, “spatial+RNA-seq”, and “spatial+RNA+sequencing”, and downloaded 579 datasets from unique GEO series (GSE) accessions. Additionally, we manually reviewed the papers in the Museum of Spatial Transcriptomics^1^ and collected 75 publicly available datasets. We also collected 115 datasets from other resources including Single Cell Portal (https://singlecell.broadinstitute.org/single_cell), 10x Genomics spatial gene expression demonstration datasets (https://support.10xgenomics.com/spatial-gene-expression/datasets), Spatial Research Lab (https://www.spatialresearch.org/resources-published-datasets/), 10x Genomics spatial publication list (https://www.10xgenomics.com/resources/publications), Amyotrophic Lateral Sclerosis Spinal Cord Atlas^2^, spatialLIBD^3^, STAR-FINDer^4^, and Brain Research through Advancing Innovative Neurotechnologies Initiative – Cell Census Network (https://biccn.org/data). Next, we removed the duplicative datasets, validated that the downloaded data used fluorescence in situ hybridization (FISH) or next-generation sequencing (NGS) based spatial transcriptomics technology, and excluded the datasets missing spatial coordinates information.

In total, we have collected 304 datasets containing 2,785 spatial transcriptomics samples from 15 different technologies and 11 species (human, mouse, rat, chicken, pig, drosophila, canine, axolotl, zebrafish, thale cress, and aspen). The human and mouse samples come from different organs (bladder, brain, breast, digits, heart, intestine, joints, kidney, lacrimal gland, liver, lung, lymph node, mouth, muscle, nasopharynx, oral mucosa, ovary, pancreas, prostate, skin, spinal cord, spleen, stomach, testis, thymus, uterus) and other specific tissues including body fat, bone, tumor cell lines, and embryonic tissues.

### Data Processing

We downloaded the count matrices and coordinate information for each dataset and applied a systematic data processing workflow to all the collected datasets. To account for the resolution and sequencing depth difference among spatial transcriptomics techniques, samples measured using different technologies were processed with corresponding quality control (QC) protocols. For 10x Visium, ST^5^, sci-Space^6^, CBSST-Seq^7^, spatial-CITE-seq^8^, and DBiT-seq^9^ datasets, we removed the capture locations with fewer than 500 unique molecular identifiers (UMIs), fewer than 500 genes, or ≥ 25% mitochondrial reads^10^. We further excluded the capture locations with a total UMI count (or a total number of genes) three standard deviations below the median^11^. Finally, we filtered out the genes that are expressed in less than five capture locations. Single-cell-resolution technologies like 10x Xenium, MERFISH^12, 13^, EEL-FISH^14^, Stereo-seq^15^, STARmap^16^, osmFISH^17^, seqFISH^12, 18,19^, and seqFISH+^20^ typically measure a smaller number of genes at lower sequencing depth. Therefore, we only performed cell QC on these datasets by removing capture locations with fewer than 500 UMIs or ≥ 25% mitochondrial reads^10^. We performed QC on Slide-seq^21, 22^ samples so that the genes with total UMI counts less than 300 were excluded^23^. Additionally, only the capture locations with total UMI counts greater than 100 and less than 25% mitochondrial reads were included^23^. **Fig. 1c** demonstrates the average number of post-QC genes per sample, and Supplementary Fig. 5 shows the average number of UMI counts and genes per capture location of samples after QC in data generated by different technologies. After QC, we normalized and transformed the raw datasets using SCTransform, a framework for the normalization and variance stabilization of molecular count data^24^. We next performed principal component analysis on normalized data and clustered the capture locations through a shared nearest neighbor approach^11^. All data processing was conducted using R v4.1.1 and Seurat V3^11^. The processed samples were stored in a standardized format including an expression table (gene by capture location) and a coordinates table (capture location by Cartesian coordinates).

### Spatial Clustering

Spatial clustering has been shown to identify spatial domains than ordinary clustering methods more accurately through jointly analyzing coordinates information and gene expression data^25^. We performed spatial clustering using STAGATE^25^, a graph attention auto-encoder framework for identifying spatial domains by learning low-dimensional latent embeddings from integrated spatial information and gene expression. STAGATE utilizes an attention mechanism to adaptively learn the similarity of neighboring spots^25^. The radius cutoff is optimized for each spatial transcriptomics technology to achieve at least five neighbors per spot. STAGATE also incorporates a cell-type-aware module by pre-clustering gene expression to enhance characterization at spatial domain boundaries^25^. Normalized counts of the top 3000 highly variable genes were used to conduct spatial clustering analysis. The number of clusters is optimized based on the Bayesian information criterion for cluster sizes between 2 and 30.

### Cell Type Deconvolution and Annotation

In order to perform cell typing, we curated reference single-cell RNA sequencing (scRNA-seq) datasets of different tissue types. We queried the GEO and identified scRNA-seq datasets with annotated cell types for each tissue type featured in SOAR. Next, we processed these datasets using an approach similar to that of the spatial transcriptomics dataset, including QC, normalization, and transformation.

In spatial transcriptomics data generated by certain technologies like 10x Visium, each capture location may contain multiple cells^26^. Therefore, in order to perform accurate cell typing, deconvoluting the cell types of each capture location is needed^26^. We performed cell type deconvolution on SOAR’s multiple-cell-resolution spatial transcriptomics datasets using BayesPrism, a Bayesian method for predicting cellular composition and cell-type-specific gene expressions in bulk RNA-seq and spatial transcriptomics data^27, 28^. To reduce batch effects, we excluded chromosomes X and Y, ribosomal, and mitochondrial genes from the analysis^27^. The genes with expression greater than 1% of the total reads in over 10% of capture locations were considered outlier genes^27^ and were also removed. To improve cell typing accuracy, we only used the cell type signature genes for deconvolution analysis^27^. The cell type markers were identified based on the differential expression analysis results on the scRNA-seq reference. The predicted cell type fractions with above 0.5 coefficient of variation were clipped to zero to reduce noise^27, 28^. We further transformed the cell-type-specific expression matrices into pseudo-cell-level by dividing the expression values by the predicted fractions of that cell type in different capture locations. The pseudo-cells with zero fraction of the considered cell type were discarded, each pseudo-cell was assigned the coordinates of its original capture location, and the data corresponding to different cell types was next combined. The created pseudo-cell-level data was used in subsequent distance-based cell-cell interaction and differential gene analysis.

For the single-cell-resolution spatial transcriptomics datasets, we performed cell type annotation using SingleR^29^, a method capable of annotating the cells in spatial transcriptomics datasets^30, 31^ based on their similarities to reference single-cell RNA sequencing (scRNA-seq) datasets with known cell types. The scRNA-seq datasets were then used as references for annotating the cell types of spatial transcriptomic capture locations of the corresponding tissue type. In particular, for non-cancer brain datasets, we adopted a heuristic-guided approach to improve the performance of cell type annotation. Two scRNA-seq datasets from the Allen Brain Map (https://portal.brain-map.org/atlases-and-data/rnaseq) were used as the references – the Human Multiple Cortical Areas SMART-seq dataset (for annotating human samples) and the Mouse Whole Cortex and Hippocampus dataset (for annotating mouse samples). Their cells were annotated as glutamatergic, GABAergic, or non-neuronal following the Common Cell Type Nomenclature (CCN)^32^. Firstly, we identified marker gene sets for each cell type and each species by performing differential gene expression analysis on the corresponding reference scRNA-seq dataset using Seurat V3^11^. Next, we used AUCell^33^ to score the activity of glutamatergic, GABAergic, and non-neuronal gene sets at each capture location based on marker gene expressions. Capture location clusters in the sample can then be classified as neuronal or non-neuronal according to the sum of AUCell scores across capture locations^33^. Finally, we used SingleR^29^ to annotate the neuronal clusters as glutamatergic or GABAergic based on a filtered version of the reference dataset that only contained neuronal cells.

### Website Development

SOAR is a comprehensive and user-friendly database that aids the exploration and analysis of spatial transcriptomics datasets. SOAR was implemented using the R Shiny framework (R v4.2.1, Shiny v1.7.1) on an Apache2 HTTP server and is compatible with smartphones and tablets. The website consists of five functional components, “Home”, “Data Browser”, “Gene & Cell Analysis”, “Drug Discovery”, “Download”, and “Help” (Supplementary Fig. 6a). The “Home” module includes an overview of SOAR, and users can search for a gene of interest in this module. In the “Data Browser” module, users can identify a sample of interest using different filters and visualize its spatial gene expressions (Supplementary Fig. 6b). Upon searching for a gene on the homepage, users will land in the “Gene & Cell Analysis” module, which enables users to evaluate the spatial variability of genes in different tissues and assess possible cell-cell interactions (Supplementary Fig. 6c). Notably, users may view the analysis results across multiple datasets using this module, allowing systematic comparisons among different samples and cell types. The “Drug Discovery” module allows users to investigate and visualize each pathological sample’s differential gene expression, protein-protein interaction, drug enrichment, and drug perturbation analysis results (Supplementary Fig. 6d). All the results and visualizations from user-performed analyses are downloadable. In the “Download” module, users can download all the curated gene expression data, coordinate data, metadata, and sample-wise analysis results. The “Help” page documents the website and includes a tutorial with step-by-step instructions for using the database. SOAR is free and open to all users at https://soar.fsm.northwestern.edu/ and there is no login requirement.

### Data Browser Module

To aid user-conducted analysis, we constructed a comprehensive data browser that is part of SOAR and contains the meta-data for all included spatial transcriptomics datasets. For each dataset, detailed information includes the hyperlink to the corresponding publication, the spatial transcriptomics technology used, and sample information including the number of samples, the species, organ, tissue, and the disease state of the sample. Furthermore, we document the average number of capture locations and genes in each sample after QC. Our data browser allows users to quickly select samples of interest to visualize spatial gene expressions, view spatial clustering and cell typing results, and perform spatial variability analysis via interactive figures and tables. All the generated figures and tables are easily downloadable to support customized and large-scale research projects.

### Gene & Cell Analysis Module

The gene search bar on the homepage of SOAR allows users to query the results of these analyses for a specific gene of interest. Upon searching for a gene, SOAR directs the user to the “Gene & Cell Analysis” tab, which subsequently prompts the user to narrow down the list of datasets by tissue type and species. In this tab, SOAR allows users to perform three types of analyses – spatial variability, neighborhood-based cell-cell interaction, and distance-based cell-cell interaction. The visualizations will be dynamically generated upon user query. All the p-values were adjusted for multiple testing using the false discovery rate (FDR) approach, and we assume statistical significance at an adjusted p-value of *q* < 0.05.

#### Spatial Variability

Studying the spatial variation of gene expression is helpful for understanding cell migration and cell-cell interaction^34, 35^. To facilitate the characterization of the functional architecture of complex tissues, we identified genes with spatial patterns of significant expression variation using SpatialDE, a statistical method for detecting spatially variable genes^35^. Spatial variability analyses were conducted across the whole tissue and in different cell types on all the samples. When evaluating cell-type-specific spatial variability, the deconvoluted expressions in individual cell types were used for multiple-cell resolution data, and the expressions in cells of the considered cell type were utilized for single-cell resolution data. Finally, we used SpatialDE to perform automatic expression histology analysis, which groups the significantly (adjusted p-value < 0.05) spatially variable genes into common spatial expression patterns^35^. This may potentially reveal histological patterns based on gene co-expression^35^.

#### Neighborhood-based Cell-Cell Interaction

Cells of different cell types may interact through cell-cell contact^36^. Spatial transcriptomics enables us to study cell-cell interactions by investigating whether a gene’s expression in one cell type appears to be promoted or inhibited when in another cell type’s neighborhood. To identify possible cell-cell interactions, we investigated whether the gene expression levels in a query cell type (*CT*_*Q*_) are influenced by its neighboring interacting cell type (*CT*_*I*_). For multiple-cell resolution datasets, each capture location was assigned the cell type with the maximum predicted fraction^37^. Denote cell type *CT*_*Q*_ capture locations as *CL*_*Q*_ and cell type *CT*_*I*_ capture locations as *CL*_*I*_. In order to evaluate neighboring interactions, we performed Wilcoxon rank-sum tests to test if genes are differentially expressed in *CL*_*Q*_ adjacent and non-adjacent to *CL*_*I*_ using the FindAllMarkers function in Seurat V3^11^. If a gene *G* is more highly expressed in *CL*_*Q*_ adjacent to *CL*_*I*_, *CL*_*I*_ may promote the expression of *G* in *CL*_*Q*_.

#### Distance-based Cell-Cell Interaction

Interactions between cells can occur beyond simple adjacency through the secretion of cytokines^36^. We evaluated the levels of cell-cell interactions through different signaling pathways in the CellChatDB database^38^ using COMMOT^39^, an optimal-transport-based approach. For multiple-cell resolution data, the pseudo-cell-level deconvoluted expression data was used. In order to characterize the cell-cell interactions over different distances, we performed COMMOT analysis using short (500 μm), medium (1,000 μm), and long (1,500 μm) distance thresholds^39^. We further identified and clustered the genes differentially expressed with respect to increased received signal through each signaling pathway using tradeSeq^40^ and COMMOT^39^. These genes may be potential downstream genes regulated by genes in the studied pathway^39^.

Both neighborhood-based and distance-based analyses can be performed when assessing cell-cell interactions. Their results can act as complementary parts and provide insight into whether the interaction happens more frequently at cell contact or over a long distance.

### Drug Discovery Module

In the “Drug Discovery” tab, users may explore the drug screening results in samples of interest using the pathological sample browser. Four analysis modules are available, including differential gene expression, protein-protein interaction, drug enrichment, and drug perturbation network, providing dynamically generated visualizations for download. All the p-values were adjusted for multiple testing using the false discovery rate (FDR) approach, and we assume statistical significance at an adjusted p-value of *q* < 0.05.

#### Differential Gene Expression

Using the FindAllMarkers function in Seurat V3^11^, we performed Wilcoxon rank-sum tests to test if genes are differentially expressed in a certain cell type compared with other cell types. For multiple-cell resolution data, the pseudo-cell-level deconvoluted expression data was used. The testing was limited to genes with at least 0.1 log fold change between two groups of cells and detected in a minimum fraction of 0.1 cells in either group.

#### Protein-Protein Interaction

For each cell type, we extracted the associated protein-protein interaction (PPI) modules using a maximum of 100 most significant differentially expressed genes and our human protein-protein interactome, which included 351,444 unique PPIs among 17,706 proteins^41–44^.

#### Drug Enrichment

To identify drug candidates with the potential for disrupting dysregulated processes in pathological samples, we conducted *in silico* drug repurposing by performing gene set enrichment analyses on the differentially expressed gene (DEG) sets associated with each cell type. We retrieved the Connectivity Map (CMap) L1000 dataset^45^ (accessed in October 2023) and analyzed perturbation profiles of 27,669 compounds associated with gene expression changes in 12,328 genes. These chemical perturbagens were treated in various cell lines and doses, resulting in a total of 145,491 perturbation profiles. Enrichment analyses were performed on the DEG sets with at least ten significantly (adjusted p-value < 0.05) up-regulated (log fold change > 0.5) or down-regulated (log fold change < −0.5) genes, and fewer than 1,000 total significant genes. We computed an enrichment score (ES) for each eligible DEG set and CMap L1000 signature using a method described previously^46–48^.

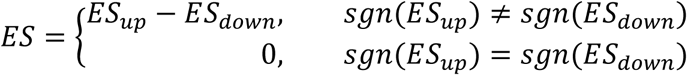

Denote the indices of the DEGs sorted by their ranks in the perturbation profiles in ascending order as *j* = 1, 2, . . . , *s*. Further denote the total number of genes in the profile as *r* and the rank of gene *j* as *V*(*j*), where 1 ≤ *V*(*j*) ≤ *r*. We calculate *ES*_*up*_ and *ES*_*down*_ for the up-regulated DEGs and down-regulated DEGs separately:

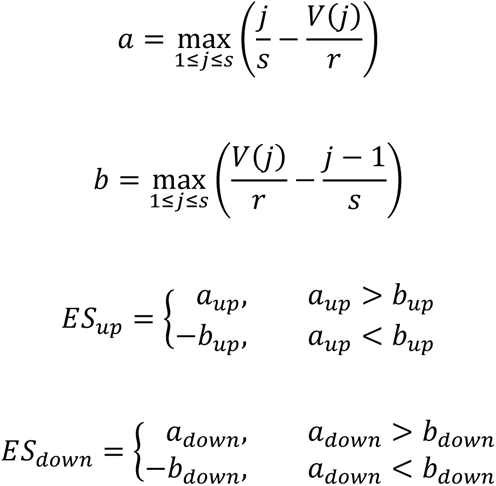

To calculate the one-tailed p-values of the ES scores, permutation tests were repeated 10,000 times using randomly generated gene lists with the same *n*(*up*) and *n*(*down*) as the DEGs. Depending on the sign of the ES score, the associated one-tailed p-value measures either the left (*ES* < 0) or right (*ES* > 0) tail. When *ES* < 0, the drug is positively related to the selected differential expression comparison and may cause a change in gene expression similar to that in the cell type of interest. On the other hand, when *ES* > 0, the drug is inversely related to the comparison and may lead to opposite gene expression patterns.

#### Drug Perturbation Network

For each set of drug enrichment analysis results, the top 100 inversely related (*ES* > 0, adjusted p-value < 0.05) and top 100 positively related (*ES* < 0, adjusted p-value < 0.05) drugs were used to generate the drug perturbation network. In the network, each node is a drug or gene, and each edge reflects an identified perturbation of the drug on the gene. The gene nodes are colored by their log fold changes in DEG analysis, and the edges are colored by the standardized perturbation scores.

## Code availability

The code we used is available at https://github.com/luoyuanlab/SOAR.

## Notes

https://soar.fsm.northwestern.edu/

### Competing Interest Statement

The authors have declared no competing interest.

### Summary of Updates

SOAR provides a comprehensive range of interactive functions for drug discovery, spatial clustering, cell typing (deconvolution and annotation), spatial expression visualization, gene spatial variability analysis, and neighborhood-based and distance-based cell-cell interaction analysis. SOAR's newly added drug discovery, spatial clustering, and cell type deconvolution functions reflect the latest developments in the field. SOAR now contains 2,785 samples from 11 species and 40 tissue types, an over 60% sample size increase from our previous submission.

