## Supplementary Information for "SOAR elucidates disease mechanisms and empowers drug discovery through spatial transcriptomics"

### Supplementary Results

#### Case Study: Drug Enrichment Analysis

In addition to sirolimus and trichostatin A, there are six other drugs, everolimus, LY294002, wortmannin, perhexiline, luminesplib, and BIBU-1361 that also showed inhibitory effects, as mediated by PI3K/Akt pathway, on DEGs of malignant cells in the case study sample.

Through competitive binding with the ATP binding site, LY294002 can inhibit the phosphorylation of PI3K, resulting in activation of GSK3 and reduced levels of the multi-drug resistance protein P-gp and anti-apoptosis proteins Bcl-2 and XIAP[1]. Activation of apoptosis through caspase-9 was also observed in carcinoma cells[2]. Combination therapy of LY294002 and idelalisib, showed reduction of reduced EMT markers, Snail and N-cadherin[3]. Wortmannin is another PI3K inhibitor with similar mechanism of action that demonstrated anti-tumor properties[4].

Luminespib is an inhibitor of the chaperon protein HSP90 required for Akt, HER3 and EGFR stability[5-7]. HSP90 inhibitor treatment leads to ubiquitination and degradation of Akt by proteasome. This disruption sensitizes tumors to chemotherapy, with higher efficacy observed in HER2 overexpressing tumors[8, 9]. Luminespib has shown potent anti-tumor activity as well as tolerability in human from phase I and II trials[10, 11].

Perhexiline is an anti-anginal CPT inhibitor. Perhexiline selectively internalized HER3 and inhibited downstream PI3K activation, thereby reducing tumor growth in breast cancer cells[12]. Perhexiline also shows inhibition toward mTORC1, as evident from the reduction of downstream RSP6 level[13]. In addition, the inhibition of fatty acid oxidation by perhexiline sensitized tumor cells to Luminespib, and the combination treatment induced apoptosis and growth arrest[14]. Lastly, perhexiline helped to overcome chemoresistance by promoting apoptosis resulted from oxidative stress and led to complete cancer regression in vivo[15, 16].

BIBU-1361 is a selective EGFR inhibitor that could prevent downstream activation of pathways involving Akt, MAPK and STAT3, thereby leading to growth arrest in tumor cells[17, 18].

### Supplementary Figures

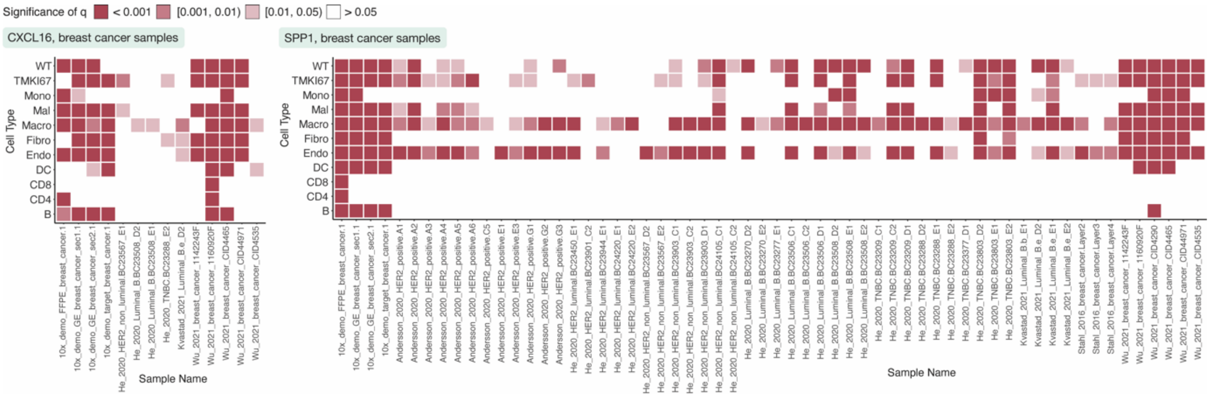


**Supplementary Fig. 1: Spatial variability of CXCL16 and SPP1 in breast cancer samples**. p-values are adjusted using the false discovery rate approach. DC, dendritic cells; Endo, endothelial cells; Fibro, fibroblasts; Macro, macrophages; Mal, malignant cells; Mono, monocytes; q, false-discovery-rate-adjusted p-value; WT, whole tissue.

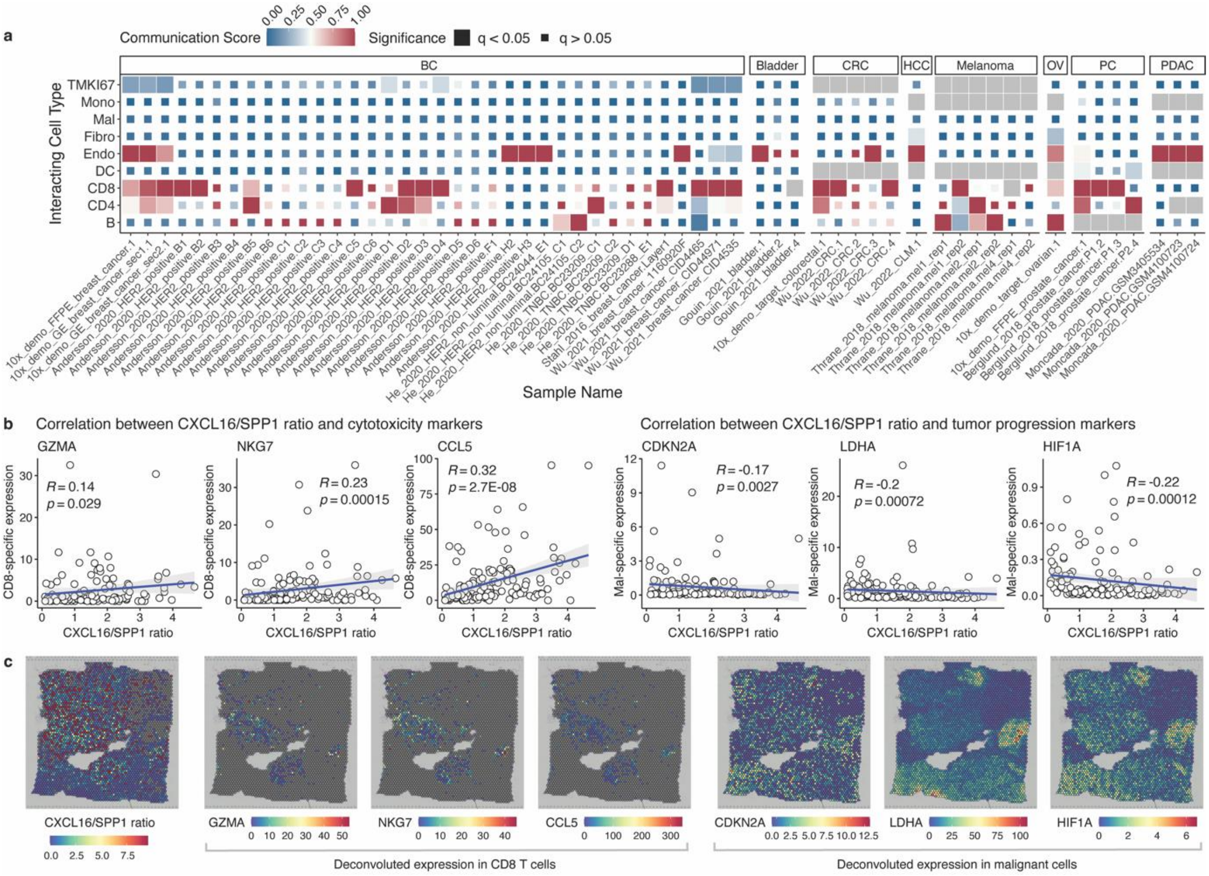


**Supplementary Fig. 2**. Supporting data for Fig. 2. **a**, Macrophages interact with T cells in cancer samples through the CXCL pathway. The tiles are colored by the communication scores, and gray color indicates that the cell type is not available in that sample. **b-c**, CXCL16/SPP1 ratio in macrophages correlates with the expression of cytotoxicity and tumor progression markers. p-values are adjusted using the false discovery rate approach. BC, breast cancer; CRC, colorectal cancer; DC, dendritic cells; Endo, endothelial cells; Fibro, fibroblasts; HCC, liver cancer; Macro, macrophages; Mal, malignant cells; Mono, monocytes; OV, ovarian cancer; PC, prostate cancer; PDAC, pancreatic ductal adenocarcinoma; q, false-discovery-rate-adjusted p-value.

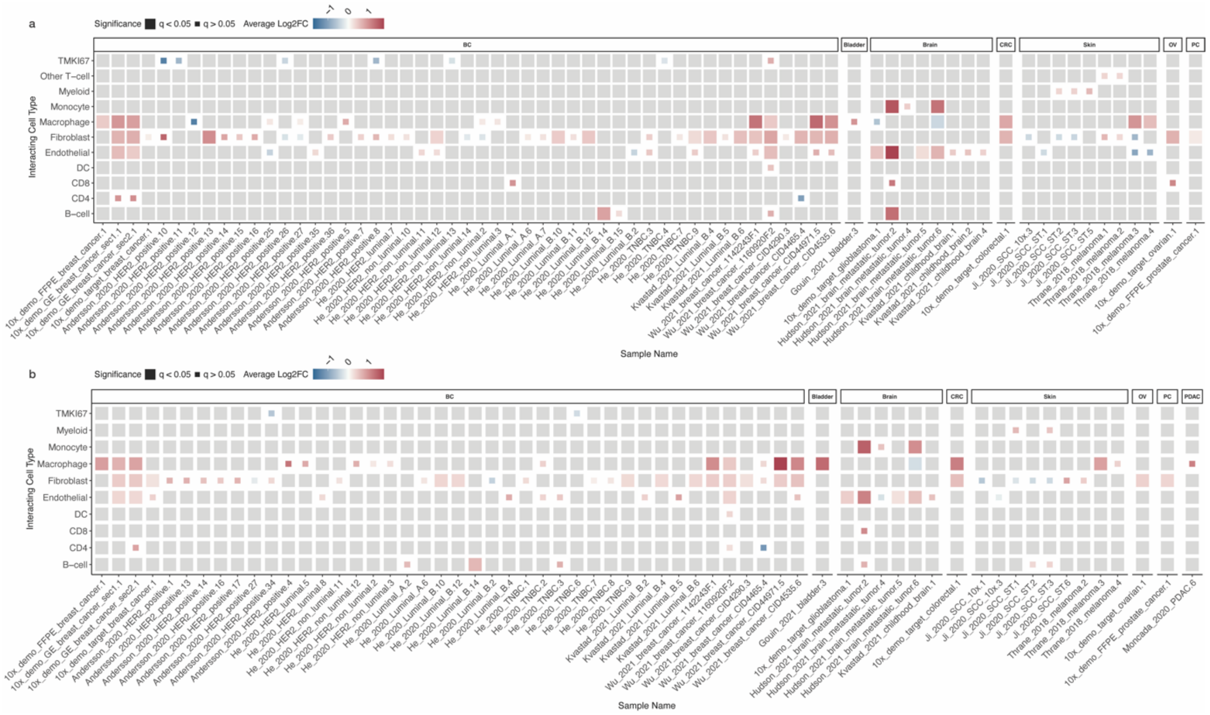


**Supplementary Fig. 3: The differential expression of tumor-promoting complement (a) C1QA and (b) C1QB between malignant capture locations adjacent and non-adjacent to other cell types.** Each tile in the heatmap is colored by the log-fold change in gene expression and sized according to its statistical significance after false discovery rate adjustment. A gray tile indicates that the cell type is unavailable in that sample. p-values are adjusted using the false discovery rate approach. BC, breast cancer; CRC, colorectal cancer; DC, dendritic cells; Log2FC, log-fold changes; OV, ovarian cancer; PC, prostate cancer; PDAC, pancreatic ductal adenocarcinoma; q, false-discovery-rate-adjusted p-value.

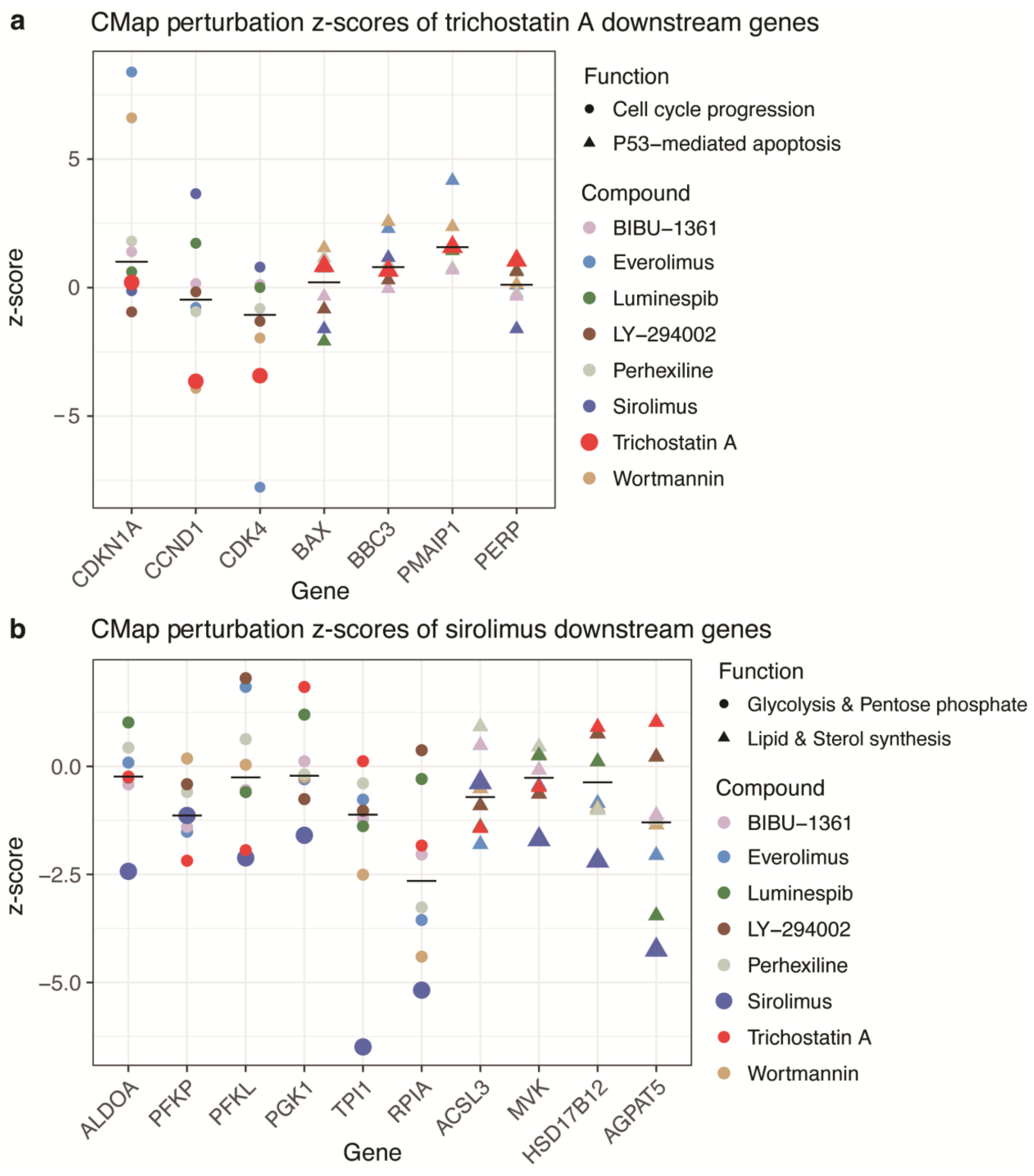


**Supplementary Fig. 4. Supporting data for Fig. 1e and Fig. 2e (drug case study) (a) CMAP shows trichostatin A downregulating cell cycle progression genes and upregulating apoptosis genes and (b) CMAP shows sirolimus downregulating energy metabolism and biosynthesis genes.** As shown from CMAP perturbation data, trichostatin A downregulates the expression of CCND1 (cyclin D1) and CDK4, which mediate G1 cycle progression and upregulates CDKN1A (p21), which is an inhibitor of the cyclin-CDK complex. In addition, trichostatin A upregulates p-53 downstream genes that contribute to apoptosis. Sirolimus downregulates the expression of genes that contribute to glycolysis, pentose phosphate, lipid, and sterol synthesis, which are all downstream targets of mTORC1. Results align with the expected mechanisms of trichostatin A as an HDAC inhibitor and sirolimus as a mTORC1 inhibitor. Perturbations of other case study drugs are also shown as a comparison. The black lines indicate median expression z scores.

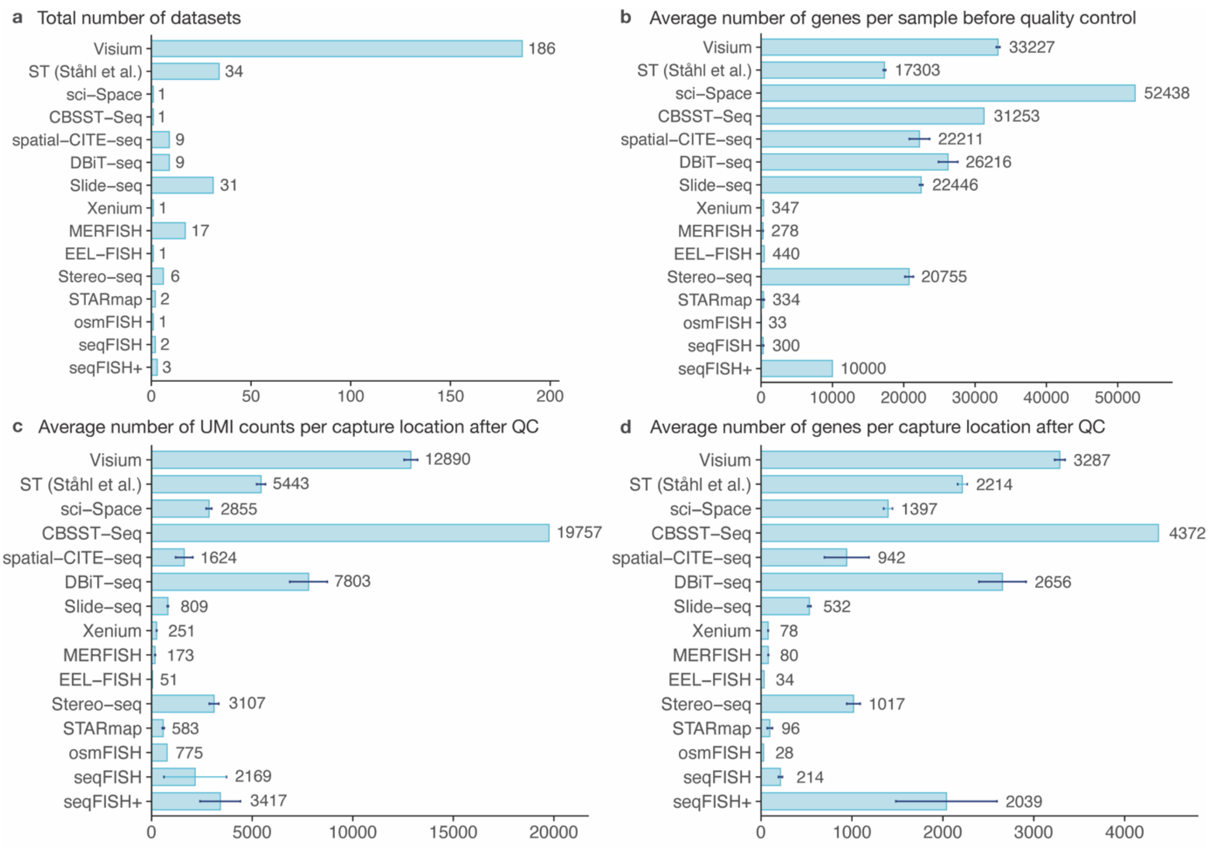


**Supplementary Fig. 5: Summary statistics of data from different spatial transcriptomics technologies**. (**a**) The number of datasets, (**b**) the average number of genes per sample before quality control, as well as the average number of (**c**) UMI counts and (**d**) genes per capture location after quality control are shown. The 95% confidence intervals for the means are plotted as error bars. QC, quality control; UMI, unique molecular identifier.

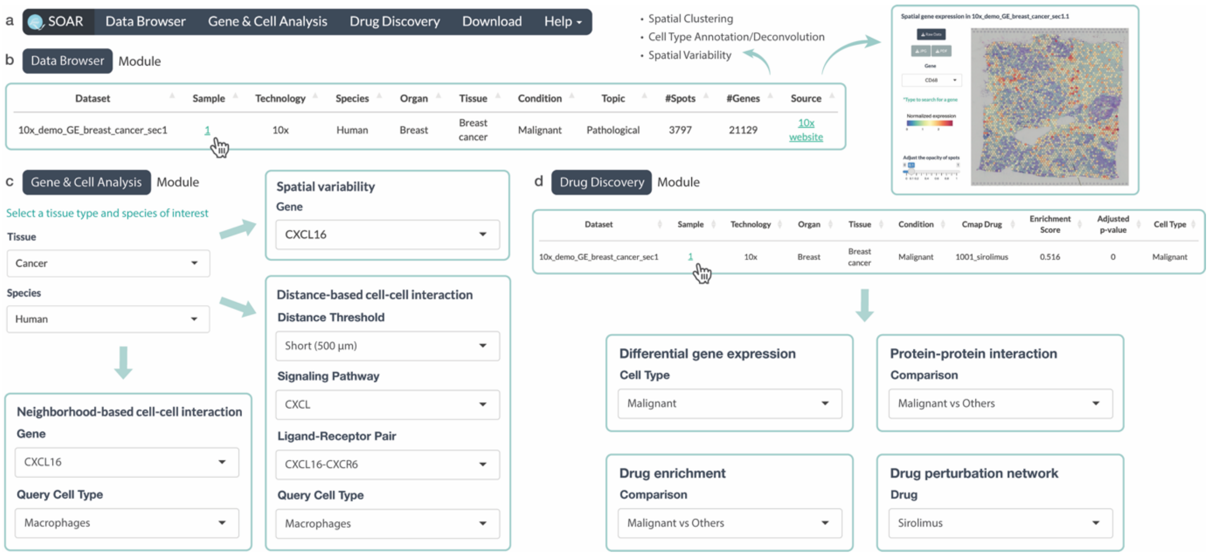


**Supplementary Fig. 6: Interactive interfaces of SOAR. a**, Main modules of SOAR, including “Data Browser”, “Gene & Cell Analysis”, “Drug Discovery”, “Download”, and “Help”. **b**, In SOAR’s “Data Browser” module, users can identify a sample of interest and visualize its spatial gene expression. **c**, In SOAR’s “Gene & Cell Analysis” module, users can first select a tissue type and species of interest. Next, users can perform spatial variability analysis or explore neighborhood-based and distance-based cell-cell interactions by interactively inputting a gene and/or query cell type. **d**, In SOAR’s “Drug Discovery” module, users can identify a pathological sample of interest and perform differential gene expression, protein-protein interaction, drug enrichment, and drug perturbation network analysis.

#### Supplementary Tables

Supplementary Table 1 is in the attached Supplementary Table 1.xlsx.

**Supplementary Table 1 Caption: Comparison of SOAR and other spatial transcriptomics resources**. SOAR contains a larger number of datasets and samples from a wider range of tissue types than existing spatial transcriptomics resources. Compared with existing spatial transcriptomics resources, SOAR has greater analysis capabilities. FISH, fluorescence in situ hybridization; NGS, next-generation sequencing.

Supplementary Table 2 is in the attached Supplementary Table 2.xlsx.

**Supplementary Table 2 Caption: Detailed information of the datasets in SOAR**. The dataset IDs are used in SOAR's "Data Browser" module. The table lists each dataset's spatial transcriptomics technology, species, and tissue type. For datasets associated with published papers and preprints, the corresponding PubMed IDs are noted, whereas, for online data collections, links to their sources are included in the table.
